# Nanopore-Based Glycan Sequencing via Fragmentation-Reassembly Strategy

**DOI:** 10.1101/2025.08.04.668447

**Authors:** Bingqing Xia, Guangda Yao, Jiamei Fan, Fangyu Wei, Jianling Tan, Pingan Li, Liuqing We, Zhaobing Gao

## Abstract

Glycan sequencing remains a major analytical challenge due to the high structural complexity, extensive branching, and isomeric diversity of glycans. Nanopore sensing has emerged as a promising single-molecule platform for glycan analysis, but existing strategies—such as hydrolysis and strand sequencing, face significant limitations when applied to highly branched or non-linear glycans. Here we report the first experimental realization of glycan assembly sequencing, a nanopore-based strategy that reconstructs full glycan structures from individually analyzed fragments. Using a biantennary complex-type N-glycan as a model, we performed controlled enzymatic digestion to generate structurally defined oligosaccharide fragments. An engineered α-HL nanopore was developed for enhanced glycan recognition, enabling the construction of a multidimensional electrical fingerprinting library. Fragment signals from unknown samples were matched to this reference dataset, and full-sequence reconstruction was achieved via fragment reassembly based on set-theoretic integration of structural candidates. This study establishes glycan assembly sequencing as a viable and modular approach to resolve complex glycoforms using nanopore technologies. The method expands the accessible sequence space beyond what is achievable with conventional strategies and provides a foundation for scalable, high-resolution glycomic analysis.

## Introduction

Nanopore technology has emerged as a promising platform for glycan analysis owing to its exceptional sensitivity to single-molecule structural variations ^1, 2^. Notably, bioengineered nanopores equipped with glycan-recognition motifs or optimized constriction geometries have enabled electrical discrimination of monosaccharides, oligosaccharides, and specific linkage isomers ^1-5^. These developments offer a high-throughput, label-free, and ultrasensitive roads of glycan detection ^6, 7^. However, direct sequencing of complex glycans using nanopores remains fundamentally challenging due to three key limitations. First, the ionic current signals generated during glycan translocation often represent a convolution of multiple interacting substructures, making it difficult to deconvolute and assign discrete events to individual monosaccharide units. Second, native glycans are typically branched, structurally flexible, and electrically neutral ^8-11^, which promotes intramolecular folding and hinders efficient translocating through the nanopore^12^. Unlike nucleic acids, glycans lack a linear backbone ^13, 14^, which allows multiple monosaccharide units to enter the pore simultaneously and significantly complicates signal interpretation. Third, the absence of a standardized “sequence reconstruction framework” severely limits the transformation of complex ionic signals into interpretable structural information. Addressing these challenges requires an integrated strategy encompassing molecular design, signal deconvolution, and structural inference.

To overcome these limitations, we previously proposed glycan assembly sequencing, a strategy that identifies and reconstructs glycans from fragments instead of relying on the translocation of the full-chain ^6^. This strategy begins with controlled hydrolysis or enzymatic cleavage to break down complex glycans into fragments compatible with nanopore sensing. The characteristic ionic current fingerprints generated from these fragments are analyzed and matched to a reference database to determine their identity and structural origin. By inferring the connectivity and conformational context among fragments, the full glycan sequence can be reassembled. Like shotgun genome sequencing and peptide assembly ^15-17^, this approach rebuilds biomolecular sequences from fragment-level data. Compared with traditional glycan analysis methods ^18-21^, glycan assembly sequencing offers several notable advantages: (1) it overcomes the physical constraints of intact glycan translocation; (2) it enables fine-grained resolution of linkage and conformational isomers through smaller, more distinguishable fragments; and (3) it facilitates integration with existing glycomics databases to streamline structural inference. As a fundamentally new sequencing paradigm, approach offers a powerful way to expand current glycomics tools and achieve high-resolution, de novo analysis of complex glycans.

In this study, we designed and experimentally validated a complete workflow for glycan assembly sequencing. Using mild enzymatic hydrolysis of biantennary complex-type N-glycans, we generated a well-defined set of oligosaccharide fragments. The fragments were detected and identified with glycan-recognition features via reference fingerprint library. By employing a fragment assembly algorithm, we successfully reconstructed the original N-glycan sequence, thereby demonstrating the first proof-of-concept of glycan assembly sequencing.

## Results

### 2.1 Design of the Assembly-Based Nanopore Glycan Sequencing Strategy

The core concept of assembly-based nanopore glycan sequencing strategy is to fragment an unknown glycan into substructures with known structure–signal associations, then decode each fragment using nanopore sensing and reconstruct the full glycan structure via a set-theoretic computational framework without requiring a reference template. As illustrated in Figure 1A, this strategy comprises six functional modules: (1) fragmentation, in which the glycan is enzymatically, chemically, or physically cleaved into structurally informative subunits; (2) nanopore sensing, where each fragment is individually translocated through an engineered nanopore to yield high signal-to-noise ionic current events; (3) feature extraction, which involves capturing multidimensional current parameters from the raw signal traces; (4) structure assembly, in which all candidate structures are integrated using set operations (e.g., intersection, union, exclusion) to reconstruct the most consistent full-length structure (Fig.1).

**Figure 1.**
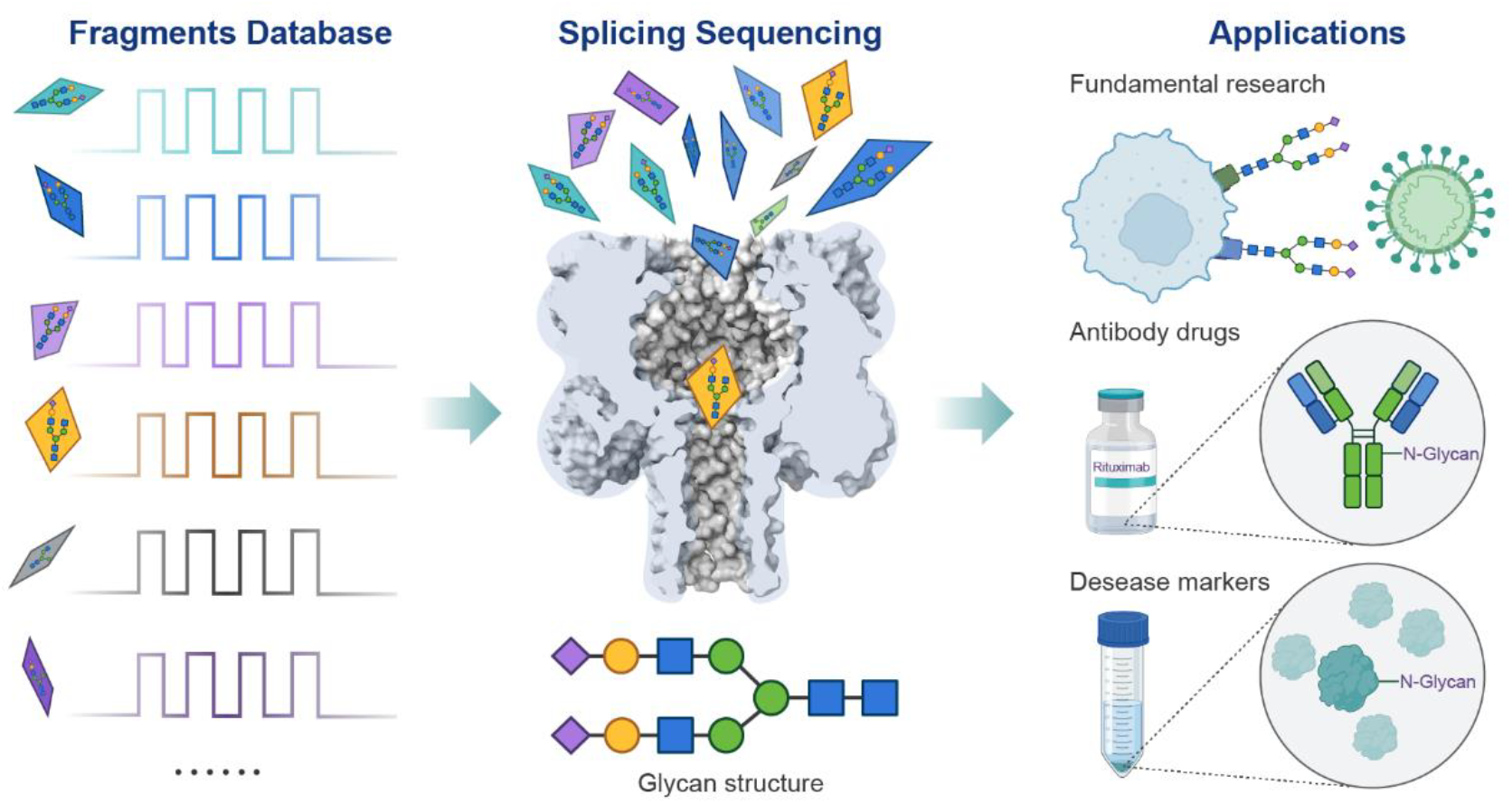
Schematic overview of the nanopore glycan splicing sequencing workflow.

### 2.2 Establishment of a Fingerprint Library for Glycan Fragments Via Engineered Nanopores

To validate the feasibility of the assembly-based sequencing strategy, we designed a panel of synthetic N-glycans with well-defined architectures and predictable fragmentation patterns as a model system. We engineered α-HL nanopores (A) to improve performance for glycan fragment detection and screened each variant for open-pore current stability under symmetric 3 M KCl (10 mM citric acid, pH 5.0) at +100 mV (Fig. 2A). To generate fragment signals for sequencing, glycans were cleaved using a combined enzymatic hydrolysis approach. Enzymes A and B were selected to generate overlapping fragments of varied lengths and isomerism (Fig. 2B). We proposed all theoretically possible fragments, labeled NF1, NF2, NF3, NF4, etc., and constructed a concept-level fingerprint library from the expected hydrolysis outcomes. Synthetic standards corresponding to key fragment candidates, including isomeric pairs (e.g., NF3 vs. NF4) and chain-length variants (e.g., NF1 vs. NF3), were prepared and analyzed using variant A nanopores (Fig. 2B). Each glycan produced frequent and clearly distinguishable blockade events (Fig. 2C). For example, fragments with sialic acid on the α1,3-arm showed shorter dwell times than those with sialic acid on the α1,6-arm (Fig. 2C). We extracted key event features, such as Dwell time and Δ*I*1/*I*0, and plotted fragment-specific fingerprint distributions (Fig. 2D-E). These fingerprints allowed reliable differentiation of the two pairs of synthetic isomers and revealed a consistent correlation between branching topology and event characteristics. Additionally, differences in chain length of the fragments were consistently resolved by the nanopore system, demonstrating its broad structural sensitivity. To confirm the single-molecule nature of the signal, we evaluated whether signal characteristics varied with analyte concentration. Over a range from 1 μM to 50 μM, the frequency of events increased proportionally, while event features such as Δ*I*1/*I*0 remained constant (Fig. 2F), consistent with single-molecule behavior. Repeated measurements of the same fragment yielded stable signal distributions with low inter-experimental variability, confirming the reproducibility and reliability of the nanopore readouts. Together, these results establish a physical and analytical foundation for assigning glycan fragments to structural identities based on electrical fingerprints, enabling downstream logical assembly.

**Figure 2.**
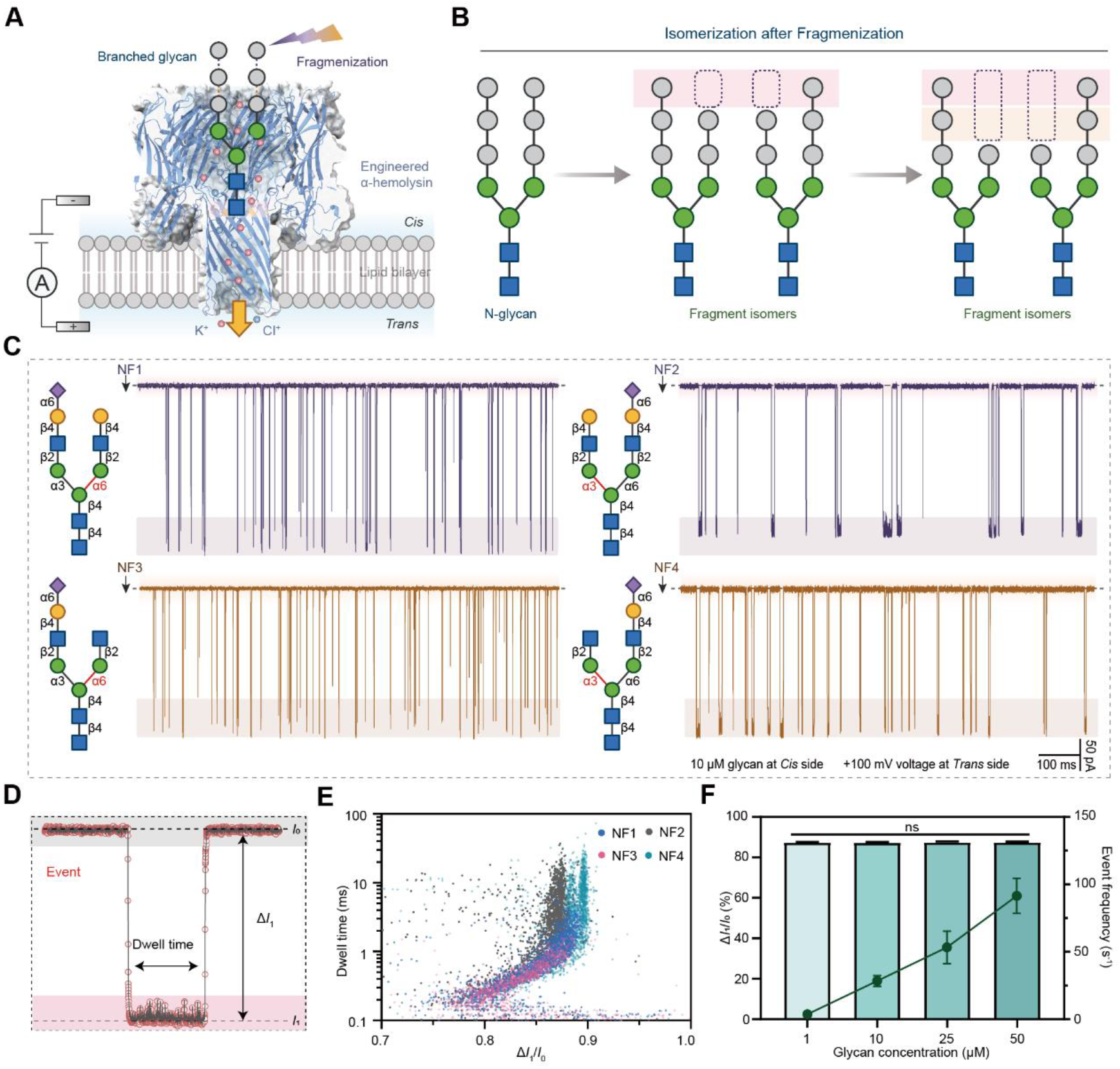
Establishment of a reference library for N-glycan fragments. (A) Principle of nanopore-based glycan detection. (B) Schematic of isomerization during glycan fragmentation via hydrolysis, where cleavage at different branches generates isomeric fragments of varying chain lengths. (C) Representative ionic current traces of four glycan fragments detected using the Site A+B/R nanopore system. (D) Characterization of current blockade events, where Δ*I*_1_ = *I*_0_ - *I*_1_ (*I*_0_: open-pore current; *I*_1_: blocked current). (E) Dwell time versus Δ*I*_1_/*I*_0_ scatter plot of NF1, NF2, NF3 and NF4. (F) Δ*I*_1_/*I*_0_ values for NF4 fragments showed no significant difference across concentration gradients, while event frequency increased with concentration. Glycans were added to the *Cis* side under a +100 mV applied voltage (*Trans* side). N ≥3

### 2.3 Automated Fragment Identification via Structure-to-Signal Mapping and Machine Learning

To enable glycan structure reconstruction without relying on structural templates, we developed a structure-to-signal lookup algorithm that integrates automated feature extraction and machine learning for real-time fragment identification based on nanopore current traces. We first established a signal processing pipeline to extract multidimensional features from each blockade event, including normalized blockade depth (Δ*I*_1_*/I*_0_), Dwell time, entry and exit slopes, and signal fluctuation standard deviation (Std). These parameters were encoded into feature vectors for downstream classification (Fig. 3A). The parameter of the fragment standards signals were used to construct a labeled training set. After denoising and normalization, 80% of events were used for training and 20% for testing. We evaluated several supervised learning models, including LDA, XGBOOST, QDA, KNN, decisiontree, SVC, NaiveBayes under cross-validation. The KNN model achieved the highest classification accuracy of 95.1%. To improve search efficiency across large fragment libraries, we implemented a two-stage retrieval process: preliminary filtering using principal component analysis (PCA), followed by fine-grained nearest-neighbor matching in the high-dimensional space. Feature importance analysis revealed that Δ*I*_1_*/I*_0_, Dwell time, and sStd were the most informative parameters. A reference dataset comprising synthetic glycan fragment standards was used to test the model, while structurally defined binary mixtures served as the test set. In a typical experiment, we prepared glycan pairs with defined structural differences, such as chain length (e.g., NF1 vs. NF3) or branching pattern (e.g., NF3 vs. NF4). Each pair was mixed at equal concentrations (2.5 μM), introduced into the *Cis* side of the mutant A nanopore, and recorded under a +100 mV holding potential for 5 minutes (Fig. 3B-C). Over 5000 single-molecule events were collected per run. Using only ∼3000 events (∼3 minutes of recording), the KNN machine learning model achieved stable and accurate fragment identification for NF1/NF3 and NF3/NF4 mixtures. (Fig. 3B-C). Overall, this automated pipeline provides a scalable, high-throughput solution for mapping single-molecule nanopore signals to glycan fragment structures, enabling robust support for downstream assembly-based sequencing.

**Figure 3.**
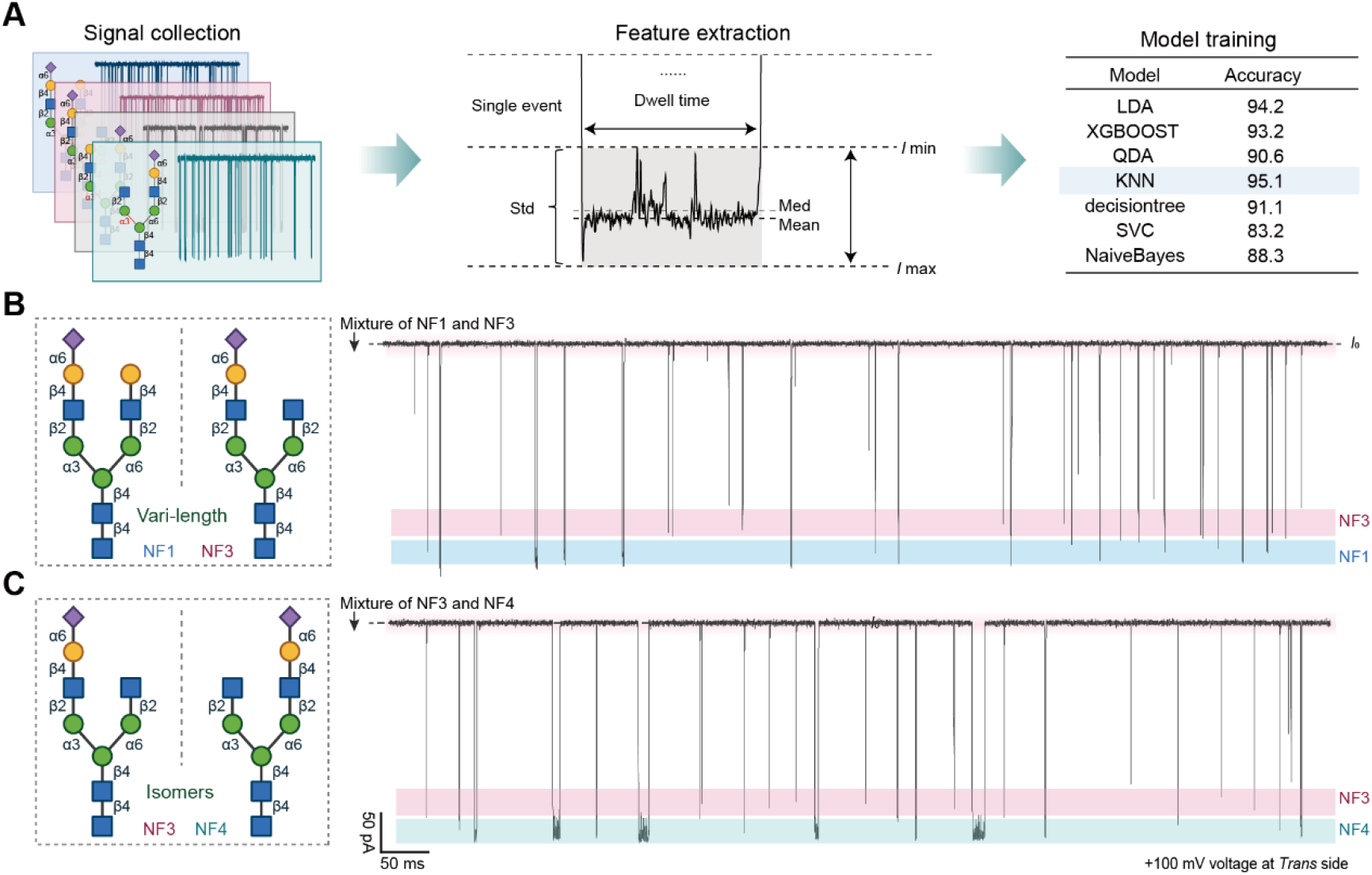
Automated Signal Extraction and Classification from Nanopore Readouts. (A) the machine learning workflow comprising current trace collection, feature extraction, model training, and structural prediction (with model accuracy comparisons shown in table). (B-C) representative current traces and structures of NF1/NF3 and NF3/NF4 mixtures

### 2.4 Implementation of Set-Theoretic Assembly and Proof-of-Concept Glycan Sequencing

To demonstrate feasibility, we used a model biantennary N-glycan (NG1) of unknown structure including its configuration. A cocktail of glycosidase A and B (0.05 mg/mL final concentration) was added to an NG solution (5mM) under controlled conditions: reaction temperature of 37 °C, pH 7.0, and timed incubations at 0, 10, 20, 30, and 60 minutes. At each time point, aliquots were taken and translocated singly through the engineered nanopore. Real-time current blockades were recorded for 5 minutes per sample to generate a time-series hydrolysate fingerprint (Fig.4A-B). Each blockade event was processed through the structure– signal lookup algorithm to assemble candidate fragment sets (Fig.4C). Parallel LC–MS analyses of the same hydrolysate supported our predicted fragment identities. After obtaining candidate structures for each fragment, we developed a set-theoretic assembly algorithm to integrate fragment-level predictions into full glycan structures. Each fragment’s Top-N candidate structures were treated as elements of a set. The algorithm applied intersection to identify shared core features, union to assemble potential branching, and difference to eliminate structural inconsistencies. This iterative process converges on the most plausible full-length structure. The workflow is illustrated in Figure 4D. We next carried out combinatorial assembly on the candidate sets. Multiple structural reconstructions were generated and scored based on internal consistency metrics, such as plausible branch connectivity and glycosidic linkage chemistry. The highest-scoring reconstruction matched the true NG1 (Fig. 4E) structure with high fidelity. This result confirms that set-theoretic operations, guided by fragment-level signal data, can reliably reconstruct complex glycan structures without a reference template.

**Figure 4.**
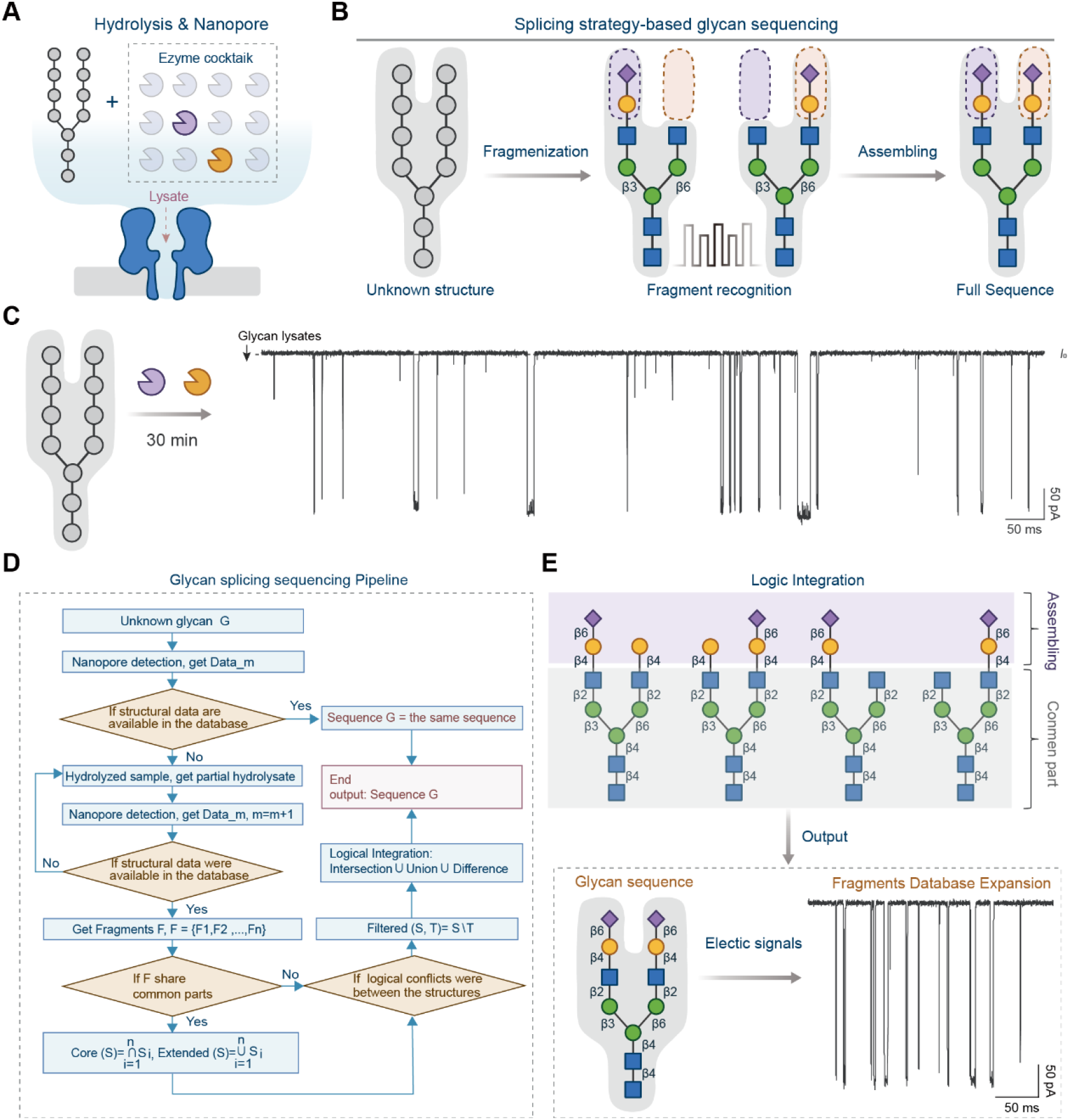
Implementation of Set-Theoretic Assembly and Proof-of-Concept Glycan Sequencing. (A) Schematic of the integrated enzymatic-nanopore detection system, with purple and orange sectors representing glycosidase A and glycosidase A, respectively. (B) Workflow of the splicing strategy-based glycan sequencing approach. (C) Representative ionic current traces from an unknown N-glycan (NG1) after 30 min enzymatic digestion by the two-glycosidase system. (D) Automated glycan assembly pipeline integrating. (E) Output of the logic integration algorithm showing the predicted N-glycan structure, with both the structural assignment and corresponding electrical signatures being incorporated into the structure-signal reference database. Glycans were added to the *Cis* side under a +100 mV applied voltage (*Trans* side). N ≥ 3

## Discussion

Here, we developed a hydrolysis-based glycan assembly sequencing strategy that integrates enzymatic fragmentation, nanopore sensing, and signal reconstruction to enable the structural decoding of complex N-glycan. Instead of translocating the whole glycan through the pore or using chemical modifications ^3, 5, 12, 22, 23^, our approach analyzes the glycan fragments to bypass topological and electrostatic difficulties. By rationally engineering the nanopore environment, we achieved highly reproducible detection of glycan fragments and constructed a multidimensional fingerprint library to represent their distinct identities. A key conceptual improvement in our approach lies in the real-time monitoring of enzymatic digestion. Rather than analyzing a static mixture of cleavage products, we continuously recorded ionic current traces from the moment of enzyme addition, capturing a dynamic and time-resolved spectrum of intermediate fragments throughout the hydrolysis process. This time-domain acquisition allowed us to observe the stochasticity of glycan degradation, whereas they would be missed in conventional endpoint analyses. All recorded fragments were computationally matched to a reference fingerprint library and assigned to their most probable substructures. Subsequently, we employed a combinatorial logic approach that examines overlapped and unique elements among candidate fragments and merges them to infer the structure of full glycan chain. This fragment-overlap strategy enhances sequencing confidence, even under conditions of partial digestion or signal noise. Together with engineered nanopores optimized for glycan recognition and machine learning classifiers for fragment identification, our workflow enables single-molecule, glycan structure reconstruction directly from enzymatic hydrolysates. While still at the proof-of-concept stage, to the best of our knowledge, our study offers the first experimental demonstration of dynamic glycan fragment capture paired with full-sequence reconstruction using a nanopore-based approach.

It worth to note, the structural assembly algorithm introduced here is conceptually original and differs fundamentally from conventional sequence reconstruction models. Traditional approaches typically require a nearly complete fragment set and apply exhaustive or heuristic alignment to reassemble the glycan, which imposes stringent demands on data coverage and often encounters exponential combinatorial complexity. These limitations become particularly pronounced when analyzing highly branched or repetitive glycan motifs ^24-26^. In contrast, our set-theory-based framework offers a mathematically rigorous and scalable alternative. By applying intersection operations to eliminate incompatible substructures and union operations to integrate redundant components, we infer the most consistent glycan backbone from incomplete data. This logic not only generalizes classical fragment-overlap strategies but also reduces the number of required observations and the size of the structural database, thereby improving tolerance to experimental variability and increasing sequencing efficiency. The formal structure of set operations further supports adaptation to diverse glycan architectures, including highly branched and compositionally heterogeneous forms. While this study remains a proof of concept, to our knowledge, it provides the first experimental demonstration of dynamic glycan fragment capture combined with full-sequence reconstruction using a nanopore-based platform, supported by an original set-theoretic assembly logic. This strategy establishes a generalizable computational foundation for glycan sequencing and may contribute to the development of more scalable glycomics technologies.

Nevertheless, glycan assembly sequencing also faces key limitations. The strategy relies on a comprehensive fragment signal library, yet the intrinsic structural diversity of natural glycans renders the construction of such a database a formidable task. Incomplete hydrolysis or signal dropout may lead to multiple plausible assembly paths, introducing ambiguity in structural interpretation. Thus, rigorous algorithmic frameworks and experimental validation are essential to ensure accurate reconstruction. These limitations highlight the need for improvements in enzymology ^27^, nanopore engineering ^28^, and computational inference to expand the applicability of this method to real-world sequencing. Despite these challenges, glycan assembly sequencing offers a fundamentally new and feasible path toward high-resolution glycan decoding. Looking forward, the modularity of this strategy facilitates its extension to other glycan classes, including O-glycans, glycolipids, and glycopeptides. Ultimately, a mature nanopore-based glycan sequencing platform capable of label-free, high-fidelity, and structurally complete readout of glycans from native biological samples holds the potential to transform the landscape of structural glycomics.

## Materials

All of glycans used in this research were synthsised by Wen Lab. The chemical reagents are listed below:

### (1) Synthetic raw materials

The key compounds used in glycan synthesis were the core N-glycan pentasaccharide and monosaccharides, including galactose and N-acetylglucosamine (GlcNAc). The mnosaccharides were purchased from BioChemSyn (www.biochemsyn.com, Shanghai, China). All other chemicals unless otherwise stated were purchased from Sigma or Carbosynth without further purification. The thin-layer chromatography (TLC) used for reaction monitoring was performed on silica gel 60 F254 plates (Merck, MA) using p-anisaldehyde sugar stain. 1H-NMR and 13C-NMR spectra were recorded on a Bruker 600-MHz NMR spectrometer (D2O as the solvent). High-resolution electrospray ionization (ESI) mass spectra were obtained using LC-MS (Thermo HPLC-Orbitrap Elite). Size Exclusion chromatography was performed using a column packed with Bio-Gel P-2 fine resins (Bio-Rad, Hercules, CA).

### (2) Protein Expression Reagents

α-hemolysin gene were chemically synthesized by BGI (Beijing Genomics Institute).

Escherichia coli BL21 (DE3) pLysS competent cells (Transgen, China); Trans8K DNA Marker (Transgen, China). tryptone (Thermo Fisher, USA), agar (Sinopharm Chemical Reagent Co., Ltd, China),agarose (Yeasen, China), Tris (Meilunbio, China), yeast extraction (Thermo Fisher, USA), ampicillin sodium salt (Yeasen, China), 2 × KeyPo Master Mix (Dye Plus) (Vazyme, China), E.Z.N.A.® Plasmid DNA Mini Kit I (Omega Bio-Tek, China), isopropyl-β-D-1-thiogalactopyranoside (IPTG, Yeasen, China), Ni Sepharose 6 Fast Flow resin (Cytiva, USA), dithiothreitol (DTT, Yeasen, China), imidazole (Yeasen, China), SDS™PAGE electrophoresis buffer powder (Yeasen, China), ethanol (Sinopharm Chemical Reagent Co., Ltd, China), Coomassie brilliant blue G250 (Yeasen, China), PageRuler™ Prestained Protein Ladder (Thermo Fisher, USA).

### (3) Materials for nanopore sensing

Potassium chloride (Sigma-Aldrich, USA), citric acid (CA) (Sigma-Aldrich, USA), Potassium hydroxide (Sinopharm Chemical Reagent Co., Ltd, China), 1,2-diphytanoyl-sn-glycero-3-phosphocholine (DPhPC) (Avanti Polar Lipids, USA), chloroform (Sinopharm Chemical Reagent Co., Ltd, China), decane (Sigma-Aldrich, USA).

## Methods

### (1) The preparation of glycans by enzymatic synthesis

The glycans were prepared by chemoenzymatic synthesis as reported previously ^24^. The N-glycans was synthesized from Core N-glycan pentasaccharide.

### (2) Nanopore protein preparation

The α-hemolysin protein was expressed and purified as described in reported research ^29^. In brief, the α-hemolysin gene with a 6× His tag at the C-terminus were inserted into the ampicillin-resistant pEASY vector by homologous recombination. Then, the plasmid was transformed into E. coli BL21(DE3) pLysS competent cells to expressed α-hemolysin protein in LB medium containing 100μg/μL ampicillin by IPTG induction. The target protein was purified by Ni-affinity chromatography. The purified α-hemolysin protein was conserved in TBS buffer (pH 8.0) and stored at -80 °C. The site-directed mutagenesis of the α-hemolysin gene was performed by PCR amplification using primers containing the desired mutation site and the template plasmid with wild-type gene.

### (3) Single-channel recordings in lipid bilayers

The single-channel recording experiments were performed as stated previously ^29^. In brief, The lipid bilayer membrane were formed across a 60 μm aperture in a PTEE film separating two chambers. 260 μL electrolyte solution (3M KCI, 10 mM citric acid, pH=5.0) were added to both chambers. The chamber connected to the ground was defined as Cis side, while the opposite chamber applied voltage was defined as Trans side. The nanopore protein were added to the Cis chamber. The analyte were added to the Cis side when the single-pore current were observed. The data were collected by nanopore instrument Cube-D2 (https://zenodo.org/records/11609574), with low-pass filtering of 5kHz and sampling rate of 50 kHz.

### (4) Features extraction and statistical analysis of current signals

The signal features were extracted from the ionic current traces using Clampfit and Python. Specifically, Clampfit was employed to extract current values from the traces and determine the start and end times of current blockade events. A Python algorithm was then utilized to calculate characteristic parameters for each glycan-induced blockade event, including standard deviation (Std), Dwell time, ΔI*1*/*I*0, mean current, median current (Med), etc. Density-based hierarchical clustering (HDBSCAN) was applied to the dataset to identify and remove noise samples.

### (5) Exoglycosidase expression and purification

Two exoglycosidases were expressed and purified in this study as described previously ^30^. The two emzyme Gene synthesis service was provided by Sangon Biotech (Shanghai, China). The gene was cloned into the pET-28a vector resulting in recombinant protein with six histidines at the protein N-terminal for further purification. The confirmed construct was subsequently transformed into E. coli BL21 (DE3) for protein expression. The His-tagged proteins were purified by using a Ni-NTA agarose column.

### (6) Glycan hydrolysis using two exoglycosidase

Hydrolysis of N-glycan: 100 μL mixture containing 5 mM of N-glycan, 0.05 μg/μL glycosidase A and B were incubated at 37°C. After the reaction was quenched by 100 °C heating for 10 min. Insoluble impurities were removed by centrifugation (10000 rpm, 5 min). The supernatant was transferred to a tube as analyte for nanopore detection.

### (7) Machine learning

After denoising and feature enhancement, a variety of supervised learning classifiers were constructed. A random seed was set to control the randomness of machine learning, ensuring the reproducibility of experimental results. Moreover, stratified K-fold cross-validation was adopted to repeatedly train and evaluate each model, with 10-fold cross-validation performed to guarantee the robustness of the results. Finally, the trained models and their corresponding preprocessing objects (standardizers, label encoders, feature orders, etc.) were saved for the prediction of subsequent unknown samples.

## Notes

The authors declare no competing financial interest.

## Acknowledgments

The authors are grateful to the Shanghai Municipal Science and Technology Major Project, the Strategic Priority Research Program of the Chinese Academy of Sciences (Grant No. XDB1360000) the Natural Science Foundation of China (Grants 92478111, 82341058), the Fund of Youth Innovation Promotion Association (Grants 2019285,and 2022077), the Shanghai Rising-Star Program (Grant 22QA1411000), the Natural Science Foundation of Shanghai (Grant 23JC1404300) and the Scientific Instrument Developing Project of the Chinese Academy of Sciences (PTYQ2024YZ0008) for financial support.

## Reference

(1) Xia, B.; Fang, J.; Ma, S.; Ma, M.; Yao, G.; Li, T.; Cheng, X.; Wen, L.; Gao, Z. Mapping the Acetylamino and Carboxyl Groups on Glycans by Engineered α-Hemolysin Nanopores. Journal of the American Chemical Society 2023, 145 (34), 18812–18824. DOI: 10.1021/jacs.3c03563.

(2) Yao, G.; Tian, Y.; Ke, W.; Fang, J.; Ma, S.; Li, T.; Cheng, X.; Xia, B.; Wen, L.; Gao, Z. Direct Identification of Complex Glycans via a Highly Sensitive Engineered Nanopore. Journal of the American Chemical Society 2024, 146 (19), 13356–13366. DOI: 10.1021/jacs.4c02081.

(3) Zhang, S.; Cao, Z.; Fan, P.; Wang, Y.; Jia, W.; Wang, L.; Wang, K.; Liu, Y.; Du, X.; Hu, C.; et al. A Nanopore-Based Saccharide Sensor. Angew Chem Int Ed Engl 2022, 61 (33), e202203769. DOI: 10.1002/anie.202203769 From NLM.

(4) Li, M.; Xiong, Y.; Cao, Y.; Zhang, C.; Li, Y.; Ning, H.; Liu, F.; Zhou, H.; Li, X.; Ye, X.; et al. Identification of tagged glycans with a protein nanopore. Nat Commun 2023, 14 (1), 1737. DOI: 10.1038/s41467-023-37348-5 From NLM.

(5) Bayat, P.; Rambaud, C.; Priem, B.; Bourderioux, M.; Bilong, M.; Poyer, S.; Pastoriza-Gallego, M.; Oukhaled, A.; Mathé, J.; Daniel, R. Comprehensive structural assignment of glycosaminoglycan oligo- and polysaccharides by protein nanopore. Nature Communications 2022, 13 (1), 5113. DOI: 10.1038/s41467-022-32800-4.

(6) Yao, G.; Ke, W.; Xia, B.; Gao, Z. Nanopore-based glycan sequencing: state of the art and future prospects. Chemical Science 2024, 15 (17), 6229-6243, 10.1039/D4SC01466A. DOI: 10.1039/D4SC01466A.

(7) Zhao, Y.; Su, Z.; Zhang, X.; Wu, D.; Wu, Y.; Li, G. Recent advances in nanopore-based analysis for carbohydrates and glycoconjugates. Anal Methods 2024, 16 (10), 1454–1467. DOI: 10.1039/d3ay02040a From NLM.

(8) Stanley, P.; Cummings, R. D. Structures Common to Different Glycans. In Essentials of Glycobiology, Varki, A., Cummings, R. D., Esko, J. D., Stanley, P., Hart, G. W., Aebi, M., Darvill, A. G., Kinoshita, T., Packer, N. H., Prestegard, J. H., et al. Eds.; Cold Spring Harbor Laboratory Press Copyright 2015-2017 by The Consortium of Glycobiology Editors, La Jolla, California. All rights reserved., 2015; pp 161–178.

(9) Griffin, M. E.; Hsieh-Wilson, L. C. Tools for mammalian glycoscience research. Cell 2022, 185 (15), 2657–2677. DOI: 10.1016/j.cell.2022.06.016 From NLM.

(10) Bertozzi, C. R.; Rabuka, D. Structural Basis of Glycan Diversity. In Essentials of Glycobiology, Varki, A., Cummings, R. D., Esko, J. D., Freeze, H. H., Stanley, P., Bertozzi, C. R., Hart, G. W., Etzler, M. E. Eds.; Cold Spring Harbor Laboratory Press Copyright © 2009, The Consortium of Glycobiology Editors, La Jolla, California., 2009.

(11) In Essentials of Glycobiology, Varki, A., Cummings, R. D., Esko, J. D., Stanley, P., Hart, G. W., Aebi, M., Mohnen, D., Kinoshita, T., Packer, N. H., Prestegard, J. H., et al. Eds.; Cold Spring Harbor Laboratory Press Copyright © 2022 by the Consortium of Glycobiology Editors, La Jolla, California. Published by Cold Spring Harbor Laboratory Press, Cold Spring Harbor, New York. All rights reserved., 2022.

(12) Lu, W.; Zhao, X.; Li, M.; Li, Y.; Zhang, C.; Xiong, Y.; Li, J.; Zhou, H.; Ye, X.; Li, X.; et al. Precise Structural Analysis of Neutral Glycans Using Aerolysin Mutant T240R Nanopore. ACS Nano 2024, 18 (19), 12412–12426. DOI: 10.1021/acsnano.4c01571 From NLM.

(13) Esmail, S.; Manolson, M. F. Advances in understanding N-glycosylation structure, function, and regulation in health and disease. Eur J Cell Biol 2021, 100 (7-8), 151186. DOI: 10.1016/j.ejcb.2021.151186 From NLM.

(14) Wilkinson, H.; Saldova, R. Current Methods for the Characterization of O-Glycans. J Proteome Res 2020, 19 (10), 3890–3905. DOI: 10.1021/acs.jproteome.0c00435 From NLM.

(15) Perret, X.; Fellay, R.; Bjourson, A. J.; Cooper, J. E.; Brenner, S.; Broughton, W. J. Subtraction hybridisation and shot-gun sequencing: a new approach to identify symbiotic loci. Nucleic Acids Res 1994, 22 (8), 1335–1341. DOI: 10.1093/nar/22.8.1335 From NLM.

(16) Zabarovsky, E. R.; Kashuba, V. I.; Pettersson, B.; Petrov, N.; Zakharyev, V.; Gizatullin, R.; Lebedeva, T.; Bannikov, V.; Pokrovskaya, E. S.; Zabarovska, V. I.; et al. Shot-gun sequencing strategy for long-range genome mapping: a pilot study. Genomics 1994, 21 (3), 495–500. DOI: 10.1006/geno.1994.1307 From NLM.

(17) Bandeira, N.; Clauser, K. R.; Pevzner, P. A. Shotgun protein sequencing: assembly of peptide tandem mass spectra from mixtures of modified proteins. Mol Cell Proteomics 2007, 6 (7), 1123–1134. DOI: 10.1074/mcp.M700001-MCP200 From NLM.

(18) Kailemia, M. J.; Ruhaak, L. R.; Lebrilla, C. B.; Amster, I. J. Oligosaccharide analysis by mass spectrometry: a review of recent developments. Anal Chem 2014, 86 (1), 196–212. DOI: 10.1021/ac403969n From NLM.

(19) Dong, X.; Huang, Y.; Cho, B. G.; Zhong, J.; Gautam, S.; Peng, W.; Williamson, S. D.; Banazadeh, A.; Torres-Ulloa, K. Y.; Mechref, Y. Advances in mass spectrometry-based glycomics. Electrophoresis 2018, 39 (24), 3063–3081. DOI: 10.1002/elps.201800273 From NLM.

(20) Grabarics, M.; Lettow, M.; Kirschbaum, C.; Greis, K.; Manz, C.; Pagel, K. Mass Spectrometry-Based Techniques to Elucidate the Sugar Code. Chem Rev 2022, 122 (8), 7840–7908. DOI: 10.1021/acs.chemrev.1c00380 From NLM.

(21) Gray, C. J.; Migas, L. G.; Barran, P. E.; Pagel, K.; Seeberger, P. H.; Eyers, C. E.; Boons, G. J.; Pohl, N. L. B.; Compagnon, I.; Widmalm, G.; et al. Advancing Solutions to the Carbohydrate Sequencing Challenge. J Am Chem Soc 2019, 141 (37), 14463–14479. DOI: 10.1021/jacs.9b06406 From NLM.

(22) Zhang, S.; Cao, Z.; Fan, P.; Sun, W.; Xiao, Y.; Zhang, P.; Wang, Y.; Huang, S. Discrimination of Disaccharide Isomers of Different Glycosidic Linkages Using a Modified MspA Nanopore. Angew Chem Int Ed Engl 2024, 63 (8), e202316766. DOI: 10.1002/anie.202316766 From NLM.

(23) Cai, Y.; Zhang, B.; Liang, L.; Wang, S.; Zhang, L.; Wang, L.; Cui, H. L.; Zhou, Y.; Wang, D. A solid-state nanopore-based single-molecule approach for label-free characterization of plant polysaccharides. Plant Commun 2021, 2 (2), 100106. DOI:10.1016/j.xplc.2020.100106 From NLM.

(24) Wei, F.; Zang, L.; Zhang, P.; Zhang, J.; Wen, L. Concise chemoenzymatic synthesis of <em>N</em>-glycans. Chem 2024, 10 (9), 2844–2860. DOI: 10.1016/j.chempr.2024.05.006 (acccessed 2025/08/03).

(25) Esmail, S.; Manolson, M. F. Advances in understanding N-glycosylation structure, function, and regulation in health and disease. European Journal of Cell Biology 2021, 100 (7), 151186. DOI: 10.1016/j.ejcb.2021.151186.

(26) Li, R.; Chen, P.; Zeng, Y. F.; Tseng, T. H.; Gannedi, V.; Krasnova, L.; Wong, C. H. Expedient Assembly of Multiantennary N-Glycans from Common N-Glycan Cores with Orthogonal Protection for the Profiling of Glycan-Binding Proteins. J Am Chem Soc 2025, 147 (15), 12937–12948. DOI: 10.1021/jacs.5c02356 From NLM.

(27) Robinson, P. K. Enzymes: principles and biotechnological applications. Essays Biochem 2015, 59, 1–41. DOI: 10.1042/bse0590001 From NLM.

(28) Samineni, L.; Acharya, B.; Behera, H.; Oh, H.; Kumar, M.; Chowdhury, R. Protein engineering of pores for separation, sensing, and sequencing. Cell Systems 2023, 14 (8), 676-691. DOI: 10.1016/j.cels.2023.07.004.

(29) Yao, G.; Xia, B.; Wei, F.; Wang, J.; Yang, Y.; Ma, S.; Ke, W.; Li, T.; Cheng, X.; Wen, L.; et al. Glycan Sequencing Based on Glycosidase-Assisted Nanopore Sensing. J Am Chem Soc 2025, 147 (2), 1721–1731. DOI: 10.1021/jacs.4c12940 From NLM.

(30) Ma, S.; Gao, J.; Tian, Y.; Wen, L. Recent progress in chemoenzymatic synthesis of human glycans. Org Biomol Chem 2024, 22 (38), 7767–7785. DOI: 10.1039/d4ob01006j From NLM.

